# DeOri 10.0: An Updated Database of Experimentally Identified Eukaryotic Replication Origins

**DOI:** 10.1101/2024.09.12.612581

**Authors:** Yu-Hao Zeng, Zhen-Ning Yin, Hao Luo, Feng Gao

**Affiliations:** Department of Physics, School of Science, Tianjin University, Tianjin 300072, China; Frontiers Science Center for Synthetic Biology and Key Laboratory of Systems Bioengineering (Ministry of Education), Tianjin University, Tianjin 300072, China; SynBio Research Platform, Collaborative Innovation Center of Chemical Science and Engineering (Tianjin), Tianjin 300072, China

**Keywords:** Replication origin, Eukaryote, Database, DNA replication

## Abstract

DNA replication is a complex and crucial biological process in eukaryotes. To facilitate the study of eukaryotic replication events, we present database of eukaryotic DNA replication origins (DeOri), a database that collects scattered data and integrates extensive sequencing data on eukaryotic DNA replication origins. With continuous updates of DeOri, the number of datasets in the new release increased from 10 to 151 and the number of sequences increased from 16,145 to 9,742,396. Besides nucleotide sequences and bed files, corresponding annotation files, such as coding sequences (CDS), mRNA, and other biological elements within replication origins, are also provided. The experimental techniques used for each dataset, as well as other statistical data, are also presented on web page. Differences in experimental methods, cell lines, and sequencing technologies have resulted in distinct replication origins, making it challenging to differentiate between cell-specific and non-specific replication. We combined multiple replication origins at the species level, scored them, and screened them. The screened regions were considered as species-conservative origins. They are integrated and presented as reference replication origins (rORIs), including *Homo sapiens*, *Gallus gallus*, *Mus musculus*, *Drosophila melanogaster*, and *Caenorhabditis elegans*. Additionally, we analyzed the distribution of relevant genomic elements associated with replication origins at the genome level, such as CpG island (CGI), transcription start site (TSS), and G-quadruplex (G4). These analysis results allow users to select the required data based on it. DeOri is available at http://tubic.tju.edu.cn/deori10/.

## Introduction

In eukaryotes, errors arising from the DNA replication process can easily lead to a series of problems, including cell damage and cancerous changes. Therefore, the mechanisms underlying DNA replication initiation and regulation need to be better understood to explore cellular changes at the molecular level. In the three domains of life, all DNA replication is initiated from replication origins [1]. However, the diversity and complexity of eukaryotic replication origins make it challenging for researchers to study them in depth and comprehensively [2, 3]. Based on numerous studies on *Saccharomyces cerevisiae* and *Xenopus laevis*, a theoretical system of eukaryotic DNA replication initiation mechanisms and regulatory systems has been preliminarily established. Although the origin of replication varies widely between phyla and species, the pre-RC mechanism is conserved in eukaryotes [4–6]. Thus, several experimental methods for replication origins have been developed, including SNS-seq (sequencing by short nascent strands), OK-seq (sequencing by Okazaki fragments), ChIP-seq (sequencing by chromatin immunoprecipitation), Repli-seq (mapping the sequences of nascent DNA replication strands throughout the six cell cycle phases), and Bubble-seq (sequencing by bubbles with restriction fragments containing replication initiation sites) [7–11]. However, the characteristics and definitions of replication origins differ slightly among species. For example, the origin of replication in budding yeast is a sequence-specific ACS sequence with a length of 100-200 bp, whereas the origin of replication in metazoans is not sequence-specific, and its length can vary from 200 bp to a 1 Mb replication domain [12, 13]. Although scientists have greatly promoted statistical analysis and data mining of replication origins, existing databases have not yet integrated massive amounts of data. Moreover, biases in samples and experimental methods, such as different protein conjugates, chromatin states and different cell stages, have resulted in different replication origins obtained by sequencing. This complexity makes it difficult for researchers to integrate and extract the required data from numerous experimental results within a short timeframe.

To address these problems, DeOri database was developed several years ago [14]. The initial version of DeOri provided 10 datasets and 16,145 sequences of replication origin. Henceforth, DeOri is constantly maintained and expanded. DeOri database has assisted experimental researchers in exploring eukaryotic replication mechanisms since its release [15], and many machine learning and statistical studies have used data from DeOri for more comprehensive analysis and prediction [16–20]. Besides, DeOri has been considered a landmark event in the study of eukaryotic replication origin [21]. In the new version of DeOri 10.0, we not only increased the number of datasets to 151 and sequences to 9,742,396, but also counted the distribution of the TSS, CpG islands, and G4 in the upstream and downstream 20kb of sequences. The genomic features within the sequence regions in each dataset were also extracted. We then provided users with distribution curves, heat maps, statistical information, and genome annotations in the sequence region (e.g., CDS, exon, mRNA, and TSS). The user-friendly search modules BLAST and JBrowse were added to DeOri 10.0, allowing users to browse, select, and compare multiple datasets more conveniently and comprehensively [22, 23]. Finally, we determined rORIs for five species with relatively conservative origins in large datasets.

## Archival resources

### Data sources and database contents

As of December 19, 2023, DeOri had collected approximately 9.7 million sequences (e.g., replication origins, replication zones, and protein-binding sites), which included 15 species, 60 cell types, and 151 datasets (Table 2). DeOri data were obtained from various experimental studies and publicly available databases. Most bed files and sequence data were obtained from Gene Expression Omnibus database (GEO) [36], and some data were obtained from other databases, such as Trypanosomatidae Database (TriTrypDB) [37] and The Encyclopedia of DNA Elements Consortium (ENCODE) [38]. Reference sequence (RefSeq) and annotations were downloaded from University of California Santa Cruz (UCSC) [39], National Center for Biotechnology Information (NCBI) [40], WormBase [41], Database of Drosophila Genes & Genomes (Flybase) [42], and The Arabidopsis Information Resource (TAIR) [43]. Based on the RefSeq and annotations, DeOri 10.0 extracted the genome features for each dataset to provide annotation files that contain the features within each sequence. The annotations included CDS, exon, mRNA, TSS, and CpG islands, which are available on the download page as a text file. DeOri also displays basic statistical information about the dataset, bed files, and FASTA files. For the data used for the analysis, TSS data were extracted from refFlat (Reference Flat File Format) files containing transcriptions in UCSC or GFF (General Feature Format) files containing reference annotations in NCBI; CpG islands data were predicted by the UCSC software cpg_lh (http://hgdownload.soe.ucsc.edu/admin/exe/linux.x86_64/cpg_lh), and G4 data were obtained from two databases: G4Bank [44] and EndoQuad [45]. Due to the differences in sequencing methods and experimental techniques in different literatures, we followed several common bioinformatics software tools with parameters used in the original literature to process the raw data. The data processing flow is shown (Figure S3). There are four main procedures: 1. Quality control; 2. RefSeq alignment; 3. Processing Bam/Sam files; 4. Peak calling. Additionally, we extracted FASTA files and genomic element annotations based on the BED files obtained from peak calling. The GC content, sequence length, and non-standard base content of sequences in the FASTA files were also calculated for users.

**Table 1.**
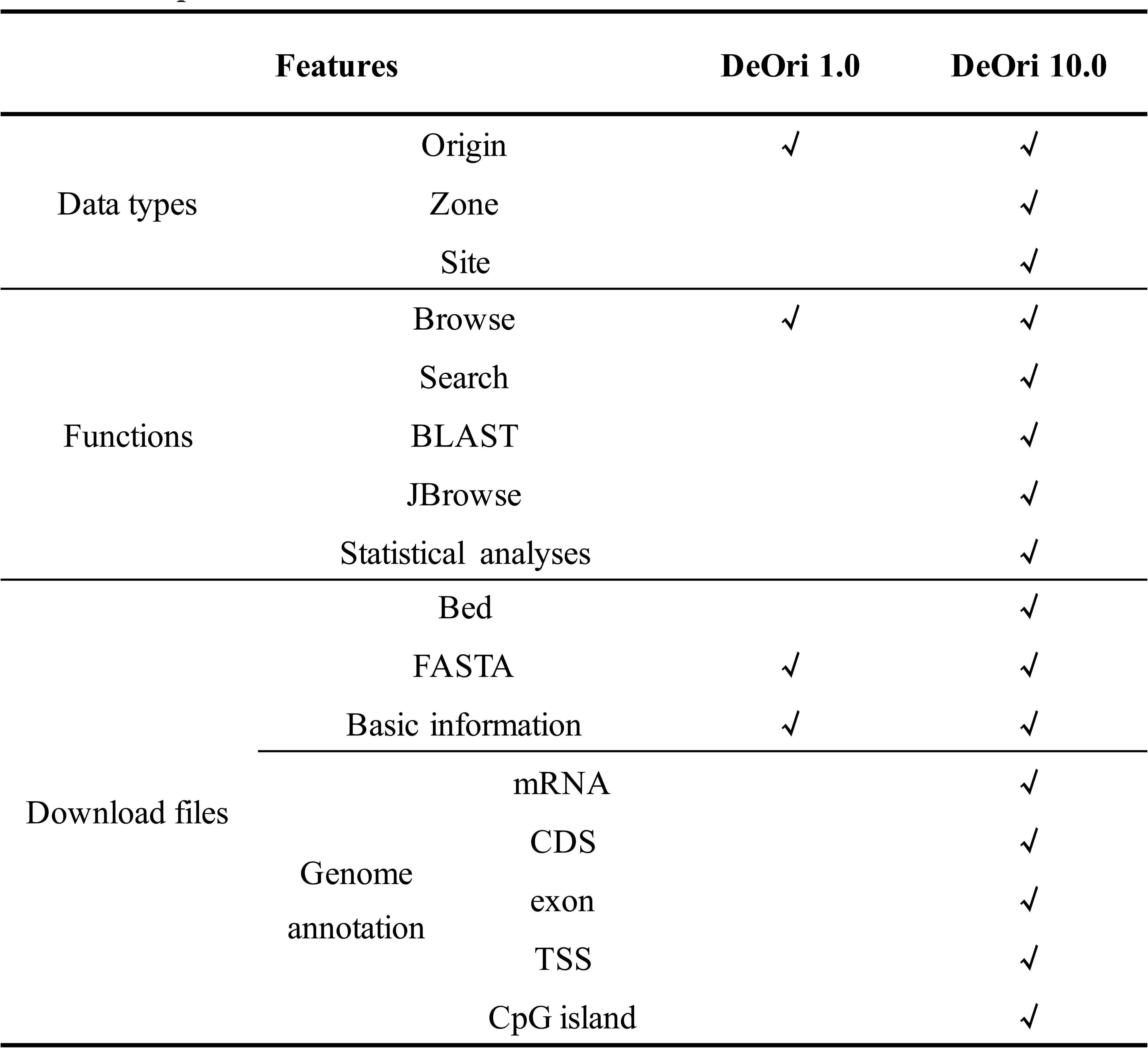
Comparison between DeOri 1.0 and DeOri 10.0.

**Table 2.**
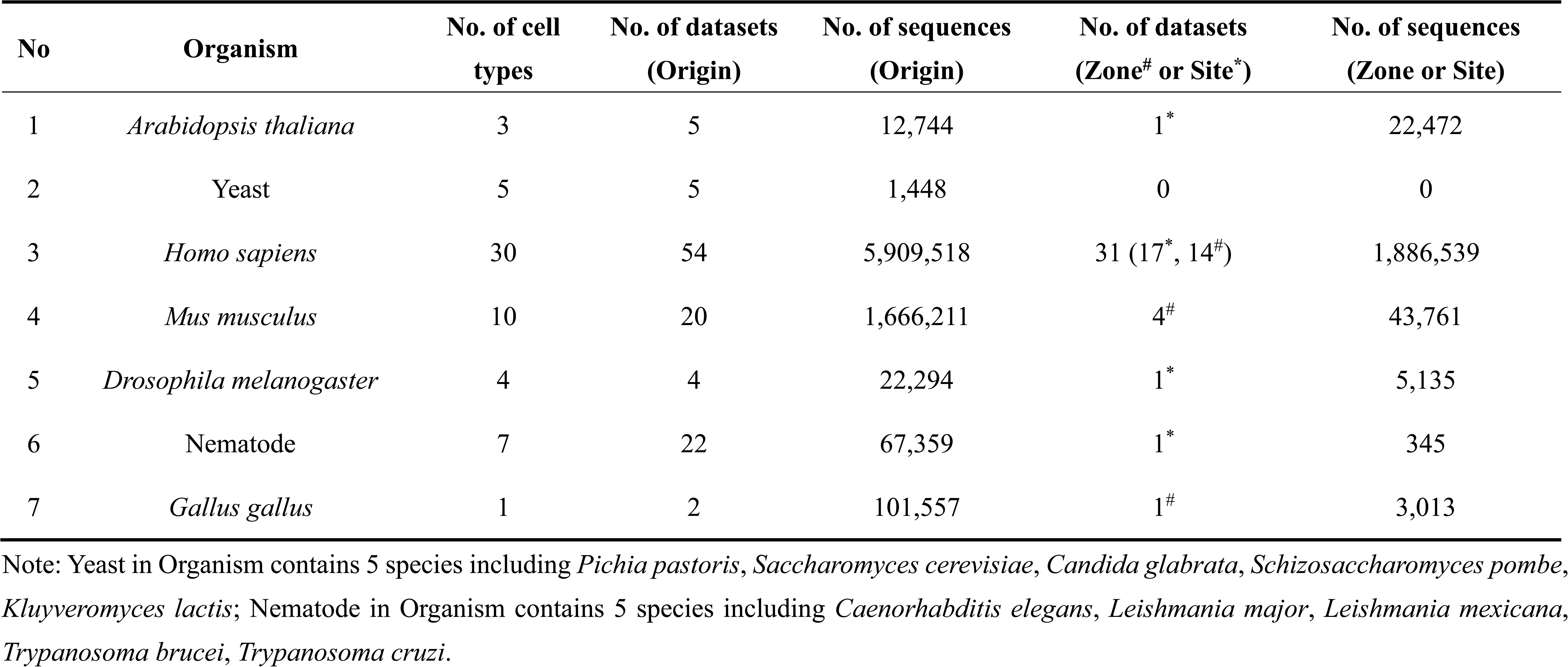
Contents of DeOri version 10.

| No | Organism | No. of cell types | No. of datasets (Origin) | No. of sequences (Origin) | No. of datasets (Zone<sup>#</sup> or Site<sup>*</sup>) | No. of sequences (Zone or Site) |
| --- | --- | --- | --- | --- | --- | --- |
| 1 | *Arabidopsis thaliana* | 3 | 5 | 12,744 | 1<sup>*</sup> | 22,472 |
| 2 | Yeast | 5 | 5 | 1,448 | 0 | 0 |
| 3 | *Homo sapiens* | 30 | 54 | 5,909,518 | 31 (17<sup>*</sup>, 14<sup>#</sup>) | 1,886,539 |
| 4 | *Mus musculus* | 10 | 20 | 1,666,211 | 4<sup>#</sup> | 43,761 |
| 5 | *Drosophila melanogaster* | 4 | 4 | 22,294 | 1<sup>*</sup> | 5,135 |
| 6 | Nematode | 7 | 22 | 67,359 | 1<sup>*</sup> | 345 |
| 7 | *Gallus gallus* | 1 | 2 | 101,557 | 1<sup>#</sup> | 3,013 |

Note: Yeast in Organism contains 5 species including *Pichia pastoris*, *Saccharomyces cerevisiae*, *Candida glabrata*, *Schizosaccharomyces pombe*, *Kluyveromyces lactis*; Nematode in Organism contains 5 species including *Caenorhabditis elegans*, *Leishmania major*, *Leishmania mexicana*, *Trypanosoma brucei*, *Trypanosoma cruzi*.

### Reference replication origins

Despite the extensive data collection in DeOri, users still face challenges in filtering and utilizing such a vast and diverse dataset. Therefore, DeOri provides users with 5 relatively conservative rORIs for eukaryotic datasets. During data preparation for rORIs, multiple datasets with different RefSeq version were transformed into the same RefSeq version using liftOver (https://genome.ucsc.edu/cgi-bin/hgLiftOver). Subsequently, the overlapping regions were scored using multiinter in bedtools [46]. For each chromosome, only regions with scores greater than or equal to 1/2 (rounded up) of the maximum overlap number were added to the reference dataset (Fig. 1). Reference replication datasets for *Homo sapiens*, *Gallus gallus*, *Mus musculus*, *Drosophila melanogaster*, and *Caenorhabditis elegans* are freely available. Based on these criteria, rORIs are regarded conservative and non-species-specific. Therefore, the rORIs are recommended for consideration when researchers give priority to sequence conservation of replication origins. In addition, rORIs can be treated as non-specific, and the origins beyond rORIs are potentially cell-specific.

**Figure 1.**
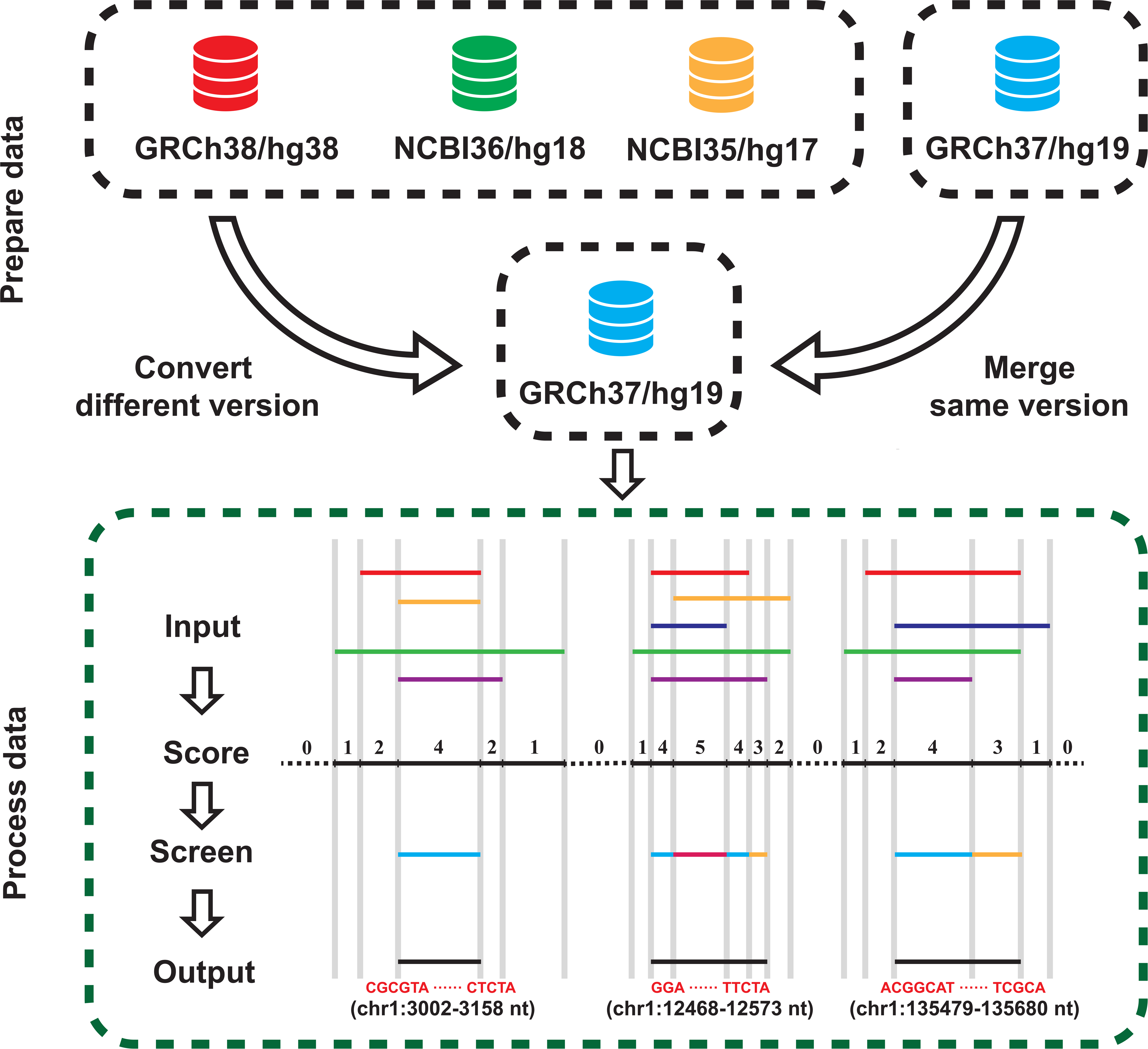
Extraction process of rORIs. Prepare data: Converting datasets of multiple different RefSeq versions in an organism into the same version; Process data: 1. Input multiple files with the same RefSeq version after conversion, and use multiinter of bedntools to score the overlapping parts of multiple datasets; 2. Get the scored file; 3. Screen the regions that are no less than 1/2 of the maximum overlapping number in the chromosome (rounded up) as the conservative replication origin; 4. Merge the adjacent fragments and output the bed file and sequences.

### Statistics on replication-related genomic elements

The statistical analysis on the distributions of genomic elements, such as TSS, CpG island, G4 and gene, was conducted at a broader scale (within the range of -20 to 20 kb from center of replication-related sequences) and a longer step size (1 kb), which would provide researchers with a landscape for further investigation. In data preparation, the files containing the location information of genomic elements are converted into bigwig format. In data calculation, the distribution matrix between each dataset and genomic elements is calculated by the computeMatrix of deepTools [47], and then the NaN value in the matrix is changed to zero. Subsequently, the columns of each matrix are averaged, and the obtained results are normalized with the scale of 0-5. Finally, the heat map is drawn based on the above calculation results. In the statistical analysis of human replication origins, datasets are merged and compared using merge and intersect of bedtools, and the correlation coefficient is calculated using jaccard of bedtools.

## Data implementation and recent updates

### Database implementation

DeOri internal framework was built using MySQL (version 8.0.35, http://www.mysql.com/) and Django (version 4.2.5, https://www.djangoproject.com/). Jinja2 (version 3.1.2), Bootstrap (version 5.3.0; https://getbootstrap.com/), EChart (version 5.4.3; https://echarts. apache.org/), jQuery (version 3.7.1, https://jquery.com/), and DataTables (version 1.13.6, https://datatables.net/) were used for page optimization and data visualization. The website was deployed and served by Nginx (version 1.20.0) and Gunicorn (version 21.2.0). Part of the picture material comes from Smart Servier Medical Art (https://smart.servier.com/).

### Updated contents and functions

Compared with the initial version, DeOri 10.0 has greatly improved and changed in data and interface (Table 1). DeOri 10.0 has been redesigned as a user-friendly and intuitive web page using the Django framework (Figure S1). There are three types of standardized data in the latest release of DeOri: Origin, Zone, and Site. The data format is slightly different, and these data type descriptions follow the original styles in the literature. Type Origin is referred to the replication origin with general definitions, whose the longest sequence is about 30kb and the average length is 0.2 kb to 5 kb in different datasets; Type Zone is referred to the region enriched replication origins or long replication initiation sequence obtained by OK-seq, whose longest sequence is about 170 kb and the average length is 20 kb to 30 kb; Type Site is referred to the region that binds to specific replication-associated protein (e.g., CDC6, MCM2-7, ORC1-2). Besides the original data, DeOri also provides datasets of reference replication origins (rORIs) in five species for users, which are believed to be relatively conservative. Taking human rORIs as an example, to verify their conservation, we extracted and screened human replication origins from datasets with no less than 10,000 sequences and compared them with human rORIs. If the rORI overlaps with the replication origins in certain dataset, this rORI is considered as overlapping rORI. The average overlap ratio is about 60%. Consequently, the percentage of overlapping rORIs to the total rORIs could indicate the conservation of rORIs and the specificity of certain datasets to some extent (Table S1). For example, although both GR00030057 and GR00030058 are immortalized HMEC cell lines, differences in the mis-expression of gene cause significant differences in their shared percentages of rORIs.

Visualizations of the dataset and sequences were presented in JBrowse. To improve the intuitiveness and convenience of DeOri, a simple home page and several modules for batch download, search, JBrowse, and BLAST are provided (Figure S2). Among these modules, JBrowse is a powerful genome browser that allows users to compare and contrast multiple datasets. It is embedded in dataset page and sequence page that demonstrates data in the JBrowse window. It can also be displayed for data browsing and comparison on a separate page. Furthermore, an example is presented on the JBrowse page.

## Data statistics

### Statistical analysis results based on DeOri 10.0

The replication origins of eukaryotes are related to particular biological elements such as CpG islands, TSS sites, and G4 structures [24, 25]. Therefore, the relevant statistical analyses were performed using DeOri 10.0. The statistical results of the relationships between these elements and the origins are presented on different pages. Users can gain an overview through the heat map on the home page (Fig. 2A) and view the data and features more intuitively on the dataset page (Fig. 2B-C). Besides, these statistical results on dataset page can be used as one of the reference factors for selecting data. Take the record GR00030030 as an example [26], which is a replication-origin dataset of the human K562 cell line. The genomic elements of TSS, CpG island, and G4 are highly concentrated within the 2 kb upstream and downstream regions from the center of replication origins in dataset GR00030030 (Fig. 2B). These statistical results not only assist users in obtaining the differences between multiple datasets, but also help to conduct further analysis.

**Figure 2.**
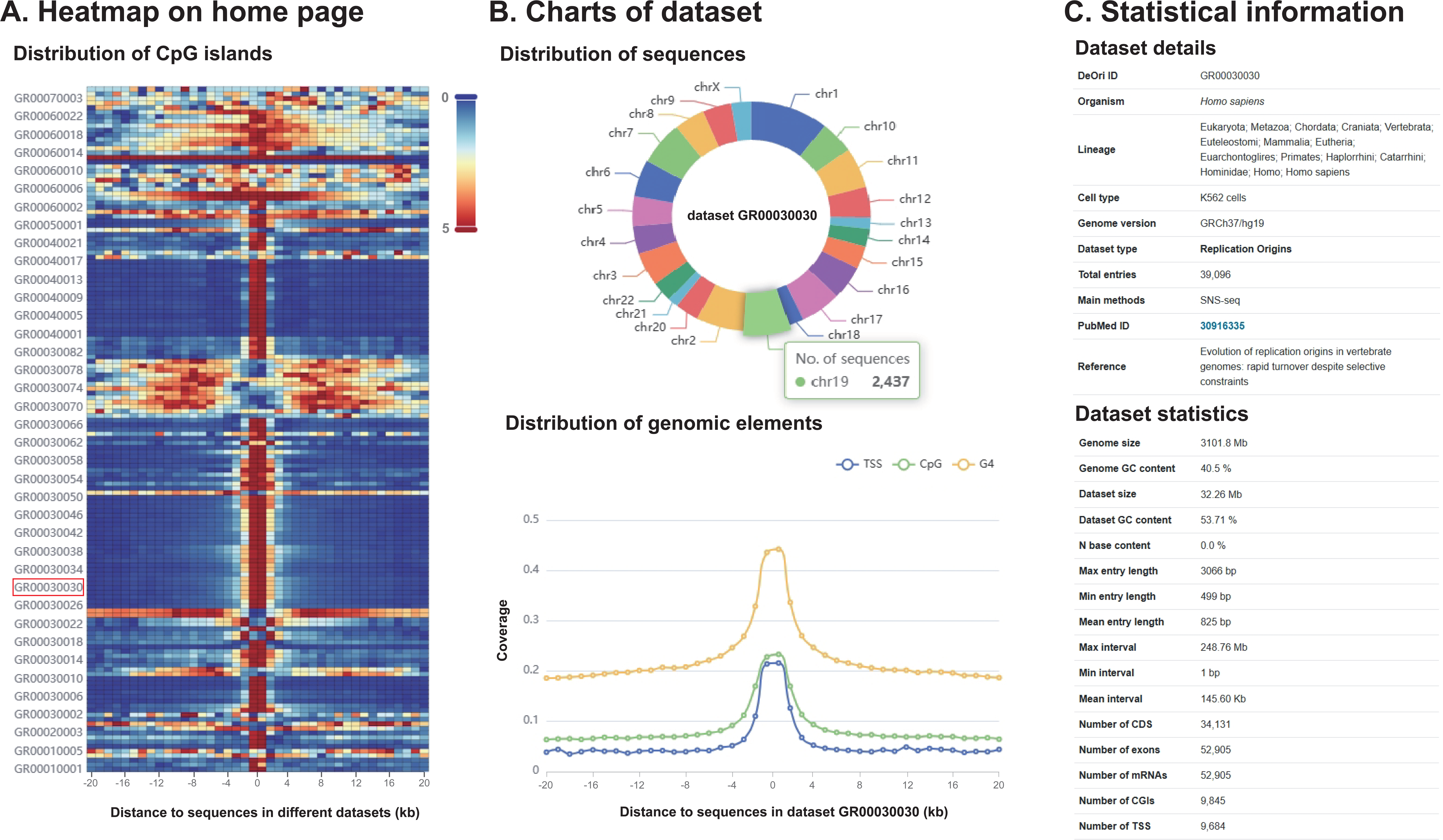
General information on datasets in DeOri. **A.** Heatmap on the home page. Comparison of distribution CpG island among different datasets. **B.** Charts of dataset. Visualization of statistics of GR00030030 dataset, such as the distribution of sequences and genomic elements. **C.** Statistical information. Basic information and statistical results on GR00030030 dataset.

### Characteristic analysis in human replication origins

To demonstrate how DeOri can be used to enhance the understanding of eukaryotic replication origins, we have conducted analysis of replication-related sequences in 85 human datasets, which have been categorized into three types: Origin, Zone and Site in DeOri 10.0 (Fig. 3A). It reveals that the GC content of type Origin is higher than that of types Site and Zone, with the latter exhibiting the lowest GC content (Fig. 3B). In order to clarify the relationship among different types, the sequences of same type were merged together, which resulted in 451,639 sequences of type Origin, 1,513,990 sequences of type Site, and 25,116 sequences of type Zone. The number of shared sequences among the three types was counted, and the Jaccard correlation coefficient was calculated, respectively (Fig. 3C). To explore the relationship between the replication-related sequences and other genomic elements, the distribution of the genomic elements, such as CpG island, G4, TSS and gene, was calculated within the range of -20 to 20 kb from the center of these sequences, respectively (Fig. 3D). Consequently, there are highly concentrated distributions of CpG island, G4, TSS and gene within the range of -2 to 2 kb for type Origin, while there are discrete distributions at both ends for type Zone. As to type Site, there are highly concentrated distributions of CpG island, G4, and TSS within the range of -2 to 2 kb, while there are no concentrated distributions of gene. Then, distributions on gene within each dataset were further calculated (Fig. 3E). Consequently, there are still highly concentrated distributions of gene in different cell lines for ORC1, 2 and CDC6 binding sites, while there is only scattered distribution at both ends for MCM2-7 binding sites (Fig. 3E). Furthermore, for all datasets, the cell lines were classified into five categories: immortal cells, differentiated cells, primary cells, multipotential stem cells, and others. It is evident that regardless of the cell lines, the majority of datasets demonstrate gene concentration within the range of -2 kb to 2kb from center of replication origins.

**Figure 3.**
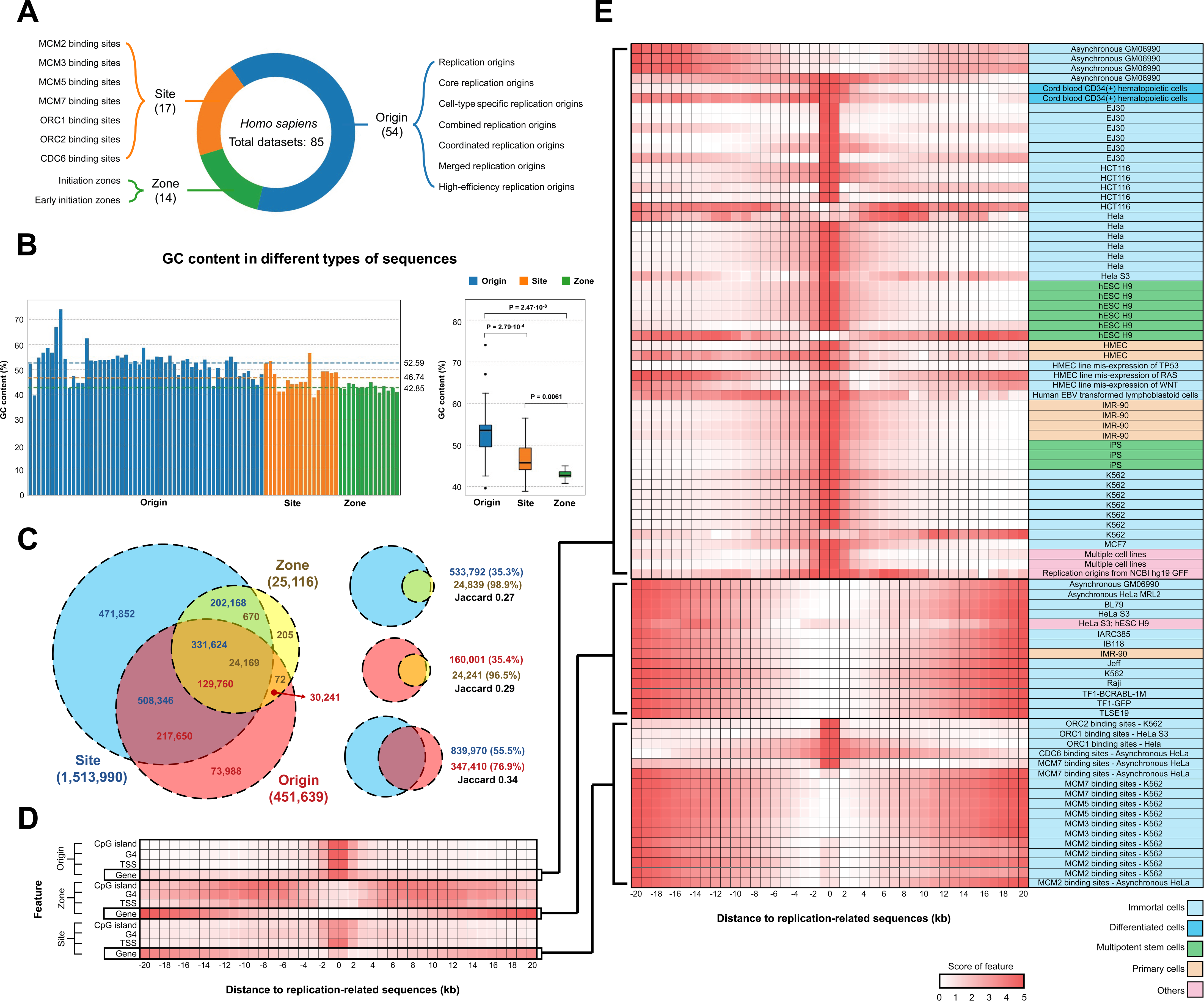
Characteristic analysis of human replication-related sequences. **A.** Statistic of human replication-related datasets in DeOri 10.0. **B.** GC content in three types of sequences. **C.** The Venn diagram of the three types of sequences. **D.** Distributions of four genomic elements around the replication-related sequences. **E.** Statistics on the distributions of gene around the replication-related sequences in each dataset.

## Conclusion and perspective

Identifying the process underlying replication origins and regulatory mechanisms has been one of a focal point in replication origin research. The proposal and validation of pre-RC and CMG mechanisms have been reported [27]. However, several questions remain unanswered. On the one hand, with the explosive growth of origin-related datasets in eukaryotes, it has been a great impetus for researchers to explore the origin characterization, firing and regulatory mechanisms of replication origins through computational analysis and prediction. However, there have been few database studies on eukaryotic origins, such as OriDB [28], which summarizes the replication origins of yeast. To fill this gap, we constructed and developed DeOri. It not only provided a large amount of experimental data, but also presented the rORIs and statistical results, which provides a lot of convenience for researchers. In practical application, researchers can compare the replication origins of different species or different cell lines based on data of DeOri, and explore their differences in sequence characteristics, chromatin distribution. For computational science, sequence features can be extracted by machine learning to predict and distinguish the replication origins. For the experimenters, they can design new experimental methods according to the experimental methods showed in DeOri. Additionally, based on the present analysis, there are highly concentrated distribution of gene around replication origins in human, offering a new analytical perspective. Subsequent analysis could focus on the specific types and functions of genes enriched in replication origins. A deeper understanding of the human replication origins may reveal differences between normal and cancer cells, potentially leading to novel treatment methods and directions.

With the rapid development of artificial intelligence and machine learning in recent years, the scientific research paradigm has been significantly promoted and changed [29]. In machine learning, many studies on replication origins have been based on the available experimental data [30–33]. For example, replication origins in bacterial genomes can be predicted using a Z-curve [34], and deep learning can be employed to identify replication origins in humans [35]. Large amounts of high-quality data are urgently required for high-quality computational science studies. For the one thing, we believe that DeOri will contribute to computational data mining and further understanding of the DNA replication process in eukaryotes. For another thing, DeOri is expected to play a crucial role in experimental and computational studies of eukaryotic replication origins. In the future, we will continue to maintain and update DeOri, and further integrate the data to eliminate redundancy of data. With the development of experimental technology and data, we are expected to add replication-related data, such as replication timing and epigenetic modification sites, in the next update to provide data services for more researchers.

## Data availability

DeOri is an open access database freely available at http://tubic.tju.edu.cn/deori10/.

## Authors’ contributions

**Yu-Hao Zeng**: Data collection, Statistical analysis, Writing - original draft, Visualization. **Zhen-Ning Yin**: Software, Writing - original draft, Visualization. **Hao Luo**: Data review, Writing - original draft, Visualization. **Feng Gao**: Conceptualization, Writing - review & editing, Supervision, Funding acquisition. All the authors have read and approved the final manuscript.

## Acknowledgements

This work was supported by the National Natural Science Foundation of China (grant numbers 32270692, 31801104, and 31571358) and the National Key Research and Development Program of China (grant number 2018YFA0903700). We thank Dr. Mei-Jing Dong for her technical help, and Professor Chun-Ting Zhang for the invaluable assistance and inspiring discussions.

## Supplementary material

**Figure S1.**
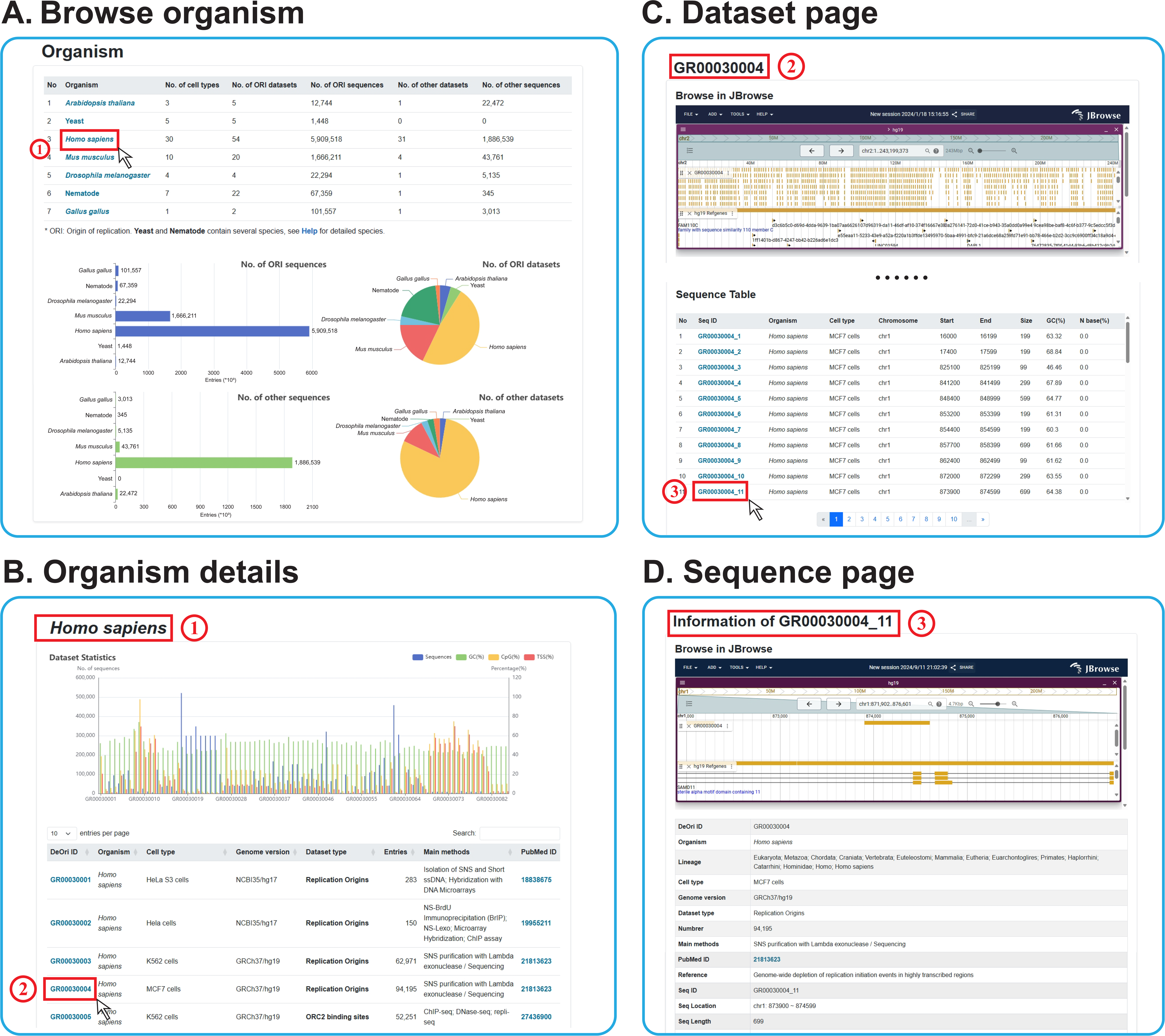
Data content of DeOri 10.0. **A.** Browse organism. Presentation of the information and statistical results on species included in DeOri. **B.** Organism details. Demonstration of the information of datasets contained in the species (e.g., cell lines, experimental methods, literature sources). **C.** Dataset page. Demonstration of the distribution of datasets, statistical information, and genome files. **D.** Sequence page. Display of details of certain sequence and the dataset on web page.

**Figure S2.**
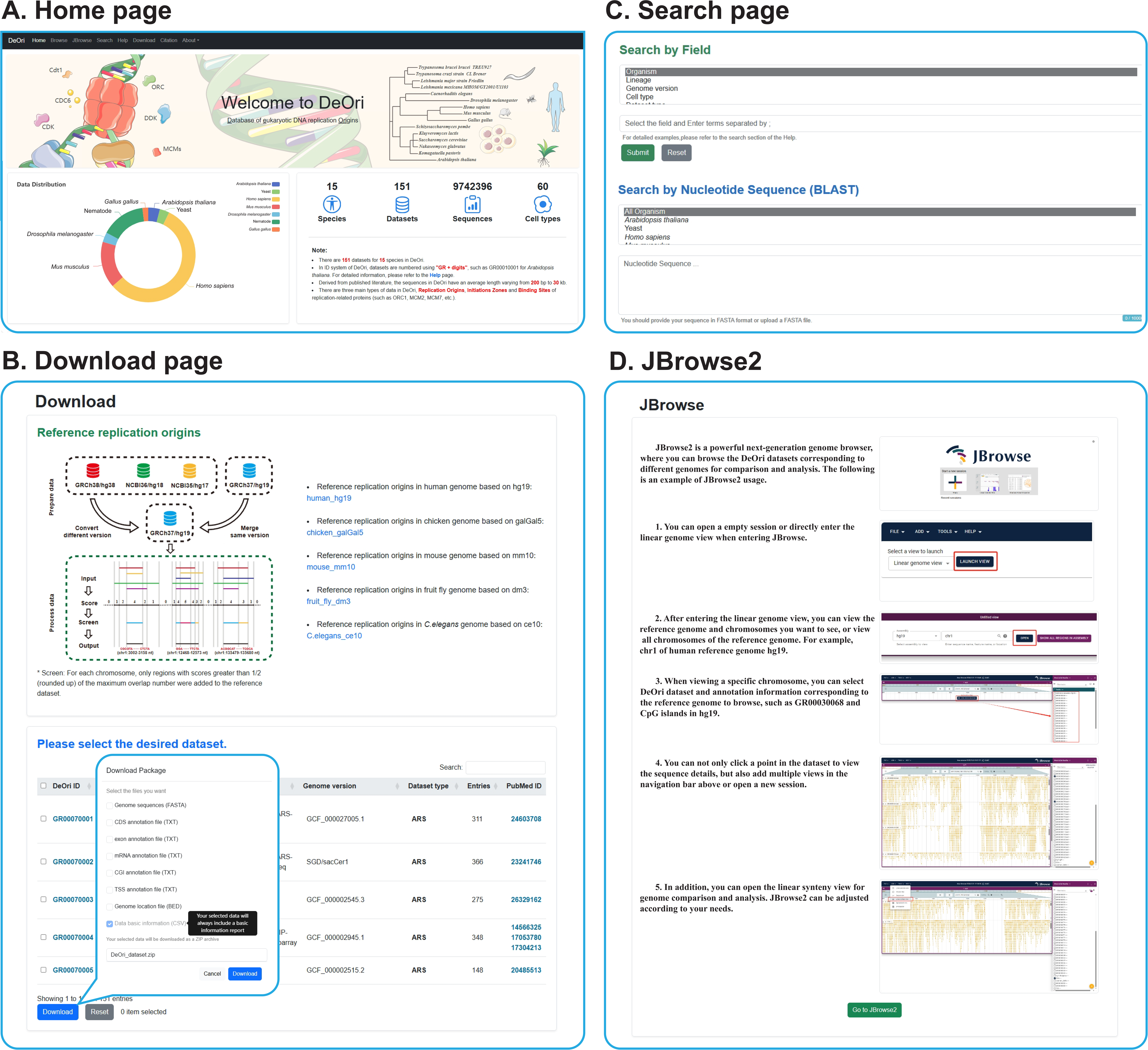
Function pages in DeOri 10.0. **A.** Home page. A basic overview of DeOri. **B.** Download pages. Presentation of datasets for data download. **C.** Search pages. Field search and BLAST based on the DeOri database. **D.** JBrowse2. An example and the entrance of JBrowse.

**Figure S3.**
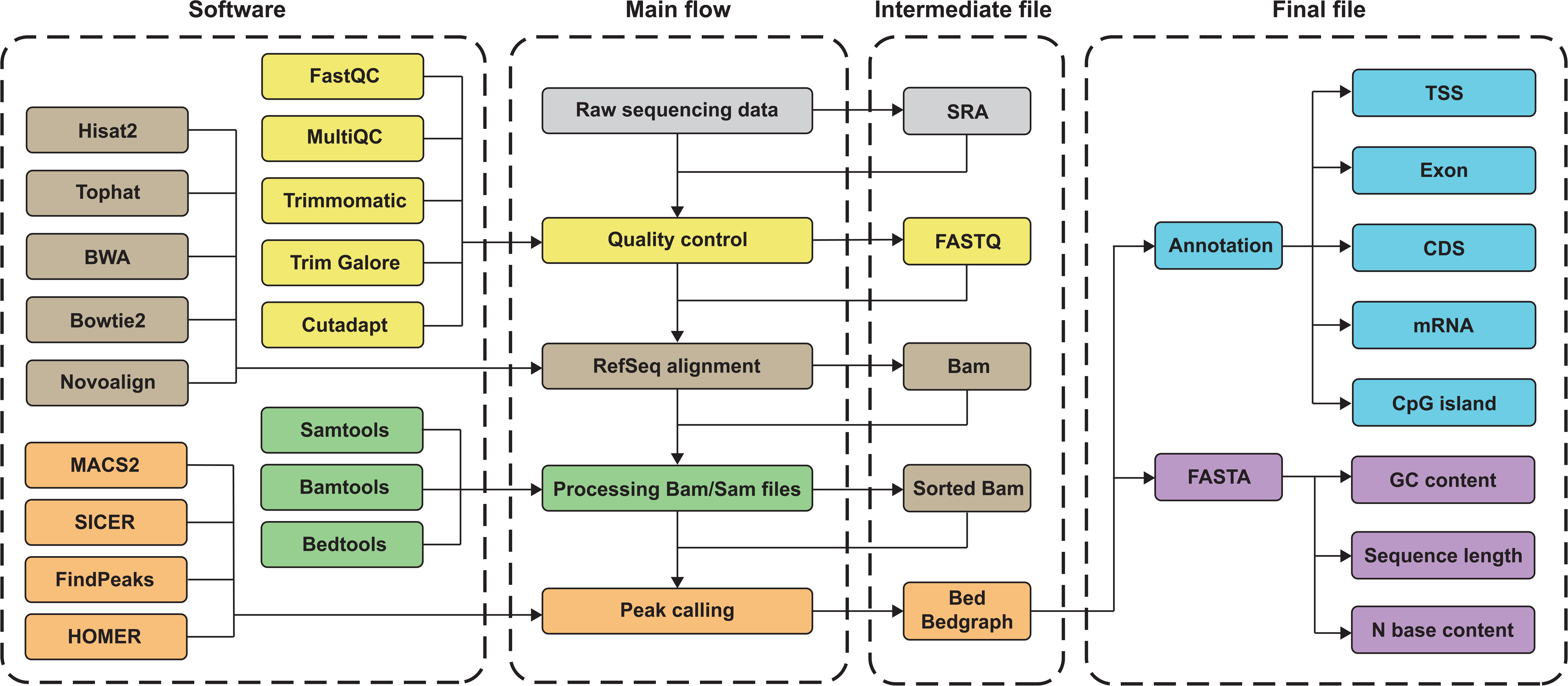
Data processing flow with related software and files.

**Table S1.** Comparison of overlap between different datasets and rORIs in human.

| DeOri ID | No. of dataset sequences | No. of rORIs | No. of overlapping rORIs | Overlap ratio* (%) |
| --- | --- | --- | --- | --- |
| GR00030003 | 62971 | 34996 | 9970 | 28.49 |
| GR00030004 | 94195 | 34996 | 13434 | 38.39 |
| GR00030005 | 52251 | 34996 | 10443 | 29.84 |
| GR00030006 | 100301 | 34996 | 22631 | 64.67 |
| GR00030007 | 119577 | 34996 | 25248 | 72.15 |
| GR00030008 | 23820 | 34996 | 15866 | 45.34 |
| GR00030009 | 10160 | 34996 | 7767 | 22.19 |
| GR00030010 | 25054 | 34996 | 11454 | 32.73 |
| GR00030013 | 123274 | 34996 | 24047 | 68.71 |
| GR00030014 | 34172 | 34996 | 17187 | 49.11 |
| GR00030015 | 39962 | 34996 | 14570 | 41.63 |
| GR00030016 | 36219 | 34996 | 13923 | 39.78 |
| GR00030027 | 41698 | 34996 | 22565 | 64.48 |
| GR00030028 | 39300 | 34996 | 20244 | 57.85 |
| GR00030029 | 39755 | 34996 | 20443 | 58.42 |
| GR00030030 | 39096 | 34996 | 20308 | 58.03 |
| GR00030031 | 39386 | 34996 | 20449 | 58.43 |
| GR00030032 | 39248 | 34996 | 20405 | 58.31 |
| GR00030033 | 115935 | 34996 | 30614 | 87.48 |
| GR00030034 | 67103 | 34996 | 29068 | 83.06 |
| GR00030035 | 84020 | 34996 | 30073 | 85.93 |
| GR00030036 | 39427 | 34996 | 20604 | 58.88 |
| GR00030037 | 166496 | 34996 | 31289 | 89.41 |
| GR00030038 | 87690 | 34996 | 23800 | 68.01 |
| GR00030039 | 143440 | 34996 | 27860 | 79.61 |
| GR00030040 | 68204 | 34996 | 27351 | 78.15 |
| GR00030041 | 151396 | 34996 | 30991 | 88.56 |
| GR00030042 | 152159 | 34996 | 30978 | 88.52 |
| GR00030043 | 85870 | 34996 | 30138 | 86.12 |
| GR00030044 | 74783 | 34996 | 29192 | 83.42 |
| GR00030045 | 128090 | 34996 | 31619 | 90.35 |
| GR00030046 | 70766 | 34996 | 27480 | 78.52 |
| GR00030047 | 155395 | 34996 | 31744 | 90.71 |
| GR00030048 | 316983 | 34996 | 29776 | 85.08 |
| GR00030049 | 63164 | 34996 | 22207 | 63.46 |
| GR00030050 | 73905 | 34996 | 20605 | 58.88 |
| GR00030052 | 95928 | 34996 | 21635 | 61.82 |
| GR00030054 | 37357 | 34996 | 11620 | 33.20 |
| GR00030056 | 90092 | 34996 | 5202 | 14.86 |
| GR00030057 | 107637 | 34996 | 22180 | 63.38 |
| GR00030058 | 110624 | 34996 | 9690 | 27.69 |
| GR00030059 | 147241 | 34996 | 25672 | 73.36 |
| GR00030060 | 184358 | 34996 | 32259 | 92.18 |
| GR00030061 | 40720 | 34996 | 14031 | 40.09 |
| GR00030065 | 124492 | 34996 | 27121 | 77.50 |
| GR00030066 | 44482 | 34996 | 19931 | 56.95 |
| GR00030067 | 78820 | 34996 | 17907 | 51.17 |
| GR00030068 | 36903 | 34996 | 3822 | 10.92 |
| Average | - | - | - | 61.16 |

Note: \* Overlap rate is calculated by dividing No. of overlapping rORIs by No. of rORIs.

